# Must epidemiologically impactful vector control interventions disrupt mosquito population structure? A case study of a cluster-randomised controlled trial

**DOI:** 10.1101/2025.04.16.649034

**Authors:** Tristan P. W. Dennis, W. Moussa Guelbeogo, Heather M. Ferguson, Steve Lindsay, Sagnon N’Fale, Patricia Pignatelli, Hilary Ranson, Antoine Sanou, Alfred Tiono, David Weetman, Mafalda Viana

**Affiliations:** School of Biodiversity, One Health, and Veterinary Medicine, University of Glasgow, Glasgow, UK; Dept. Vector Biology, Liverpool School of Tropical Medicine, Liverpool, UK; Centre National de Recherche et de Formation sur le Paludisme, Ouagadougou, Burkina Faso; Université Joseph Ki-Zerbo, Ouagadougou, Burkina Faso; Department of Biosciences, Durham University, Durham, UK; Université Yembila Abdoulaye Toguyeni, Fada N’Gourma, Burkina Faso; Groupe de Recherche Action en Santé, Burkina Faso

**Keywords:** Vector populations, genetics, disease vectors, malaria, cluster-randomised controlled trial, epidemiology

## Abstract

Large epidemiological impacts resulting from disease vector control interventions are typically associated with significant disruption of vector populations. While vector density is a frequently measured response, impacts on demography and connectivity are suspected but rarely quantified. We analysed low-coverage whole-genome sequence data of 893 *Anopheles gambiae* mosquitoes collected between 2014-2015 during a cluster-randomized control trial (cRCT) in Burkina Faso to compare a pyrethroid-only net (ITN) with a pyrethroid-pyriproxyfen (ITN-PPF) net. Despite reductions of clinical malaria by 12%, and vector density by 22% in the ITN-PPF arm, we found no significant changes in *An. gambiae* population genetic structure or diversity. We found remarkably low population differentiation, and a lack of discernible clustering by treatment, time, or space. Nucleotide diversity and inbreeding coefficient remained stable between treatments, and genome-wide scans showed no putative signatures of selection between trial arms. These results show that ITN-PPF did not alter *An. gambiae* genetic structure, possibly due to large, vagile populations in West Africa. More widely, this is first evidence that epidemiologically meaningful reductions in vector density may not impact genetic diversity or connectivity, and challenges what constitutes adequate vector control in large populations.

## Introduction

Vector control is the cornerstone of the public health response to many vector-borne diseases (VBD)^1^. Effectiveness is typically assessed by the epidemiological impacts vector control generates, such as reduction in the prevalence or incidence of infection^2,3^. These epidemiological impacts are tied to reductions in human exposure to vector populations^4^, which can arise through reductions in vector population size, adult mosquito survival or infection rates. Notable examples of vector control strategies that have large epidemiological impacts include the deployment of spatial repellents that reduced arboviral infection by 34.1% and *Aedes* mosquito abundance by 38.6% in Peru^5^ and release of *Wolbachia*-infected *Aedes aegypti* that resulted in a 78% reduction in dengue incidence in a trial in Indonesia^6^. For malaria, the deadliest VBD, responsible for over 600,000 deaths annually^7^, the effects of vector control have been profound, with a 40% reduction in clinical disease incidence across sub-Saharan Africa between 2000 and 2015^8^ as a result of mass insecticide treated bednet (ITN) distribution. Successful malaria vector control has been associated with broader changes in mosquito vector demography and ecology, including changes in biting and resting behaviour^9,10^, species composition^11^ and the emergence of insecticide resistance^12^. For example, vector control has reduced *Anopheles gambiae* populations over time in East Africa, leading to *An. arabiensis* and *An. funestus* replacing it as the dominant vectors of malaria^13,14^. These examples suggest that that large epidemiological impacts are linked to, and may typically require, significant perturbation of mosquito vector populations, including reducing population size^15^ or fragmenting population structure^16^. However, the demographic effects of interventions on vector populations are difficult to measure directly and hence remain elusive and rarely recorded.

Insect populations, such as butterflies^17–20^ and gypsy moths^21,22^, have provided notable case-studies of how populations are regulated, maintained and structured as well as how population fragmentation can lead to population declines by breaking connections among subpopulations^18,23–25^. However, the impact of vector control on mosquito population dynamics, in particular population structure, remains poorly understood. Vector control may change mosquito population structure by reducing local population size below the threshold for persistence, and in doing so compromise metapopulation viability, or population fitness by driving down diversity. In contrast, in the presence of migration or strong density-dependent regulation, local mosquito populations may persist even in the face of substantial reduction due to vector control^26,27^. It is thus important to understand the mechanisms underlying population resilience, and the magnitude of reduction required to achieve an epidemiological benefit, to improve the design of vector control programmes.

Changes in population size and structure leave genomic signatures in populations^28^. Reductions in nucleotide diversity (π and θ*w*) and increases in individual inbreeding (*F*_*IS*_) are often associated with population decline, which in turn is linked to a greater risk of stochastic extinction^23,24,29^. Furthermore, changes in rates of dispersal between subpopulations may affect metapopulation viability. Inference of close-kin relationships using genetic techniques (close-kin mark-recapture, or CKMR) has been used to quantify *Aedes* mosquito dispersal^30,31^, but are currently untested in estimating *Anopheles* taxa. Population genetics has also been used previously to estimate vector control impacts. Notable examples include a study in Kilifi, Kenya, which documented genetic signatures of population decline in *An. gambiae* following large-scale ITN deployment from 2006. Reductions in nucleotide diversity (∼5 % lower π and ∼15 % lower θ*w*), and a deficit in the number of low-frequency sites (negative Tajima’s D) were accompanied by changes in vector species composition, whereby the presence of the primary malaria vector species *An. gambiae sensu stricto* declined from 79% of *An. gambiae* species complex samples between 1997-1998 to an undetectable level in 2007-2008^32^. In a separate study in western Kenya, a shift in *An. gambiae* from 85% to 1% of collected adults in Kenya between 1999 and 2010 was coincident with high ITN coverage^13^, with genomic signatures of population decline such as increased inbreeding, decreased genetic diversity, and allele-frequency shifts, evident in samples sequenced in a separate study^33^. There are also reported instances where vector control was not associated with significant signatures of population disruption. In Cameroon, ITN implementation was associated with insignificant impacts on *An. arabiensis* genetic diversity and population structure^34^, though no clinical or entomological endpoints were reported with these data. Evidence of changes to population structure in the face of successful vector control is currently limited and variable.

Significant epidemiological impacts in cluster-randomised controlled trials (cRCT) of vector control are often accompanied by declines in vector population densities. However, the impact on mosquito population structure and the extent to which epidemiological impact requires population disruption, remains unclear. This lack of clarity could stem from the lack, to our knowledge, of published studies with systematic comparisons between treatment versus control groups that also incorporate population genetic estimations of changes in structuring or diversity. This knowledge gap can be addressed by applying population genomic techniques to dense vector sampling from clinical trials, especially field cRCTs comparing control and treatment groups. Low-coverage whole-genome-sequencing (WGS) approaches (e.g. a depth-of-coverage of <5X) are particularly attractive for capture of accurate population allele frequencies and fine-scale population structure from large samples sizes at much reduced per-sample cost than medium-full (e.g. 10-30X) coverage approaches^35,36^.

Validation of new vector control strategies, such as gene-drive approaches, spatial repellents and tools using novel insecticide classes with different entomological modes of action often require evaluation in cRCTs to determine their public health value. While standard guidelines are in place for measurement of entomological and epidemiological endpoints^37^, methodologies to understand how these interventions impact vector populations more holistically are missing. Population genetic analysis of vectors coupled with cRCTs can provide more information on the effect of vector control on mosquito population dynamics and in turn the extent to which epidemiological outcomes are coupled with substantial demographic impacts on vector populations.

Here, we use the case study of the AvecNet cRCT performed in Burkina Faso^38^, in which a standard pyrethroid insecticide-treated net (Olyset ITN) was compared with a novel net that combined pyrethroid with an insect growth regulator, pyriproxyfen (Olyset DUO net also known as ITN-PPF). The ITN-PPF reduced the incidence of clinical malaria in children by 12%, and vector numbers by 22%, compared to the standard ITN^38^. Our objective was to analyse population genetic data from *An. gambiae* samples collected during and after the trial to assess the impacts on: 1) the spatial genetic structure; 2) genetic diversity and inbreeding (as general, although imperfect^39^ proxies for population size); 3) putative signatures of selection in the genomes of mosquitoes collected from treated vs control clusters (see **Methods** for more detailed justification). We provide a framework to detect changes in population structure in the context of field trials, to give insight into the impacts of control on fragmentation and elimination of vector populations.

## Methods

### Study design and entomological sample collection

Briefly, the AvecNet stepwise cRCT was a two-arm trial undertaken in 2014–15 in Banfora, Burkina Faso^38^. In this area, malaria is endemic and largely transmitted by species of the morphologically-identical *An. gambiae s*.*l*. complex (*An. arabiensis, An. gambiae* and *An. coluzzii*). The mosquito population, and consequently malaria incidence, is highly seasonal with peaks in abundance generally concentrated during the rainy season, which occurs between July and November.

The cRCT aimed to compare the effectiveness against clinical malaria of the Olyset DUO net (hereafter ITN-PPF or treated), which in addition to the pyrethroid permethrin contains pyriproxyfen, an insect growth regulator, to the standard Olyset pyrethroid-only ITN (hereafter ITN or control), impregnated with permethrin only. The trial included 81 consenting villages that were grouped into 40 clusters, each including an average of 50 children under 5 years old. At the start of the malaria transmission season in June 2014, five randomly-chosen clusters were provided with ITN-PPF bednets, while the remaining clusters were given ITN bednets. Five clusters were randomly chosen for further ITN-PPF delivery each month from July-Nov 2014 so that by the end of 2014, each trial arm had an equal number of clusters. In 2015, ITN-PPF were distributed in a similar fashion starting in June 2015. By the end of the trial in December 2015, all clusters had the ITN-PPF nets (**See Figure 1** for more information on net and cluster distribution). During the cRCT, mosquitoes were collected using CDC Light Traps in six randomly-chosen households per cluster, every four weeks between May-December in 2014 and May-November in 2015. All mosquitoes were identified using microscopy, and a random subset of approximate 30% of *An. gambiae s*.*l*. was typed by PCR.^38^ Full details of the study protocol and collections are available in Tiono et al^38^.

**Figure 1.**
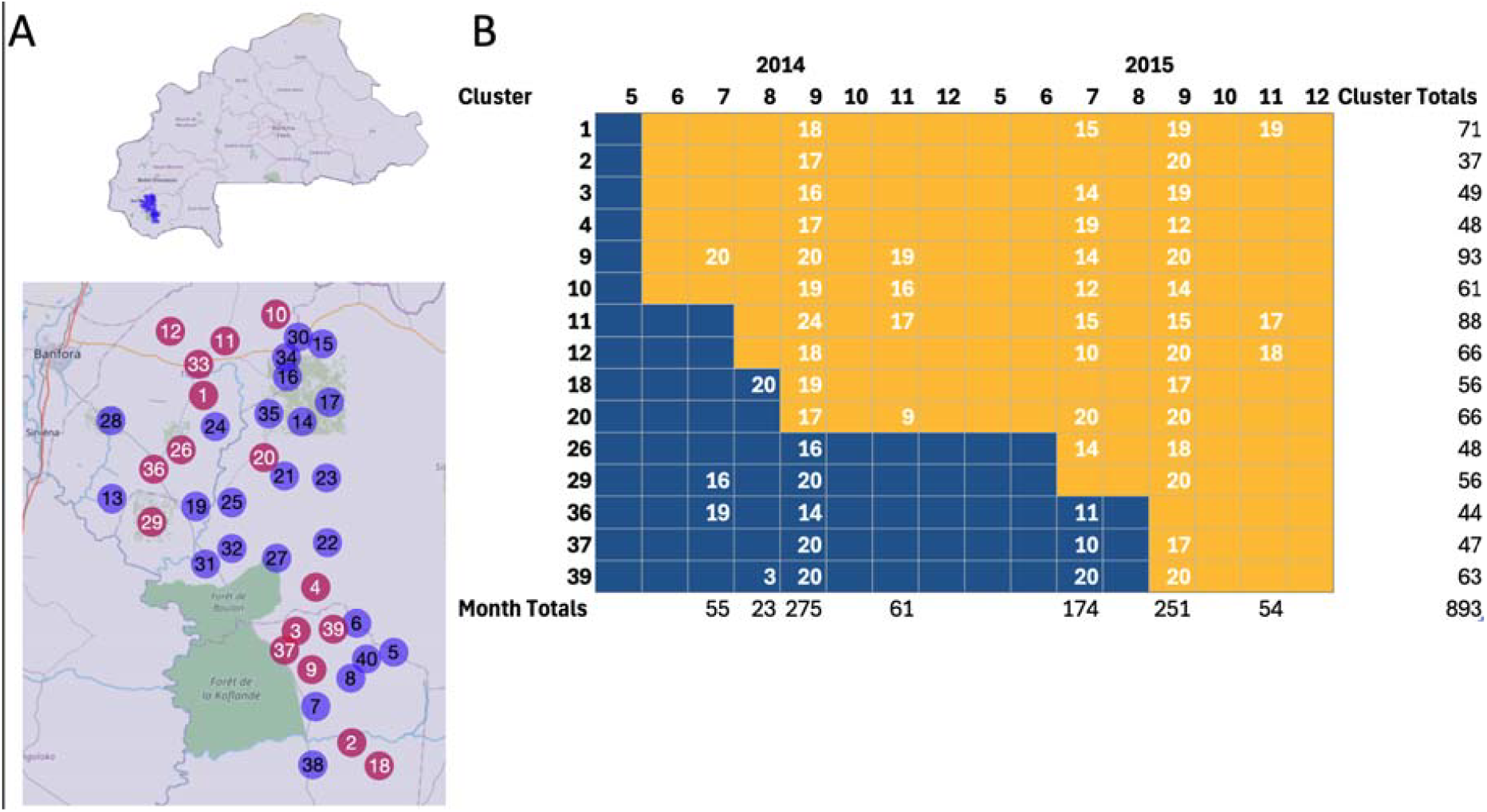
Clinical trial and collection setup. A) Map of Burkina Faso highlighting the Banfora region (top panel) and a zoom in on the centroid of the 40 clusters enrolled in the cRCT (bottom panel). Maroon circles indicate clusters from which mosquitoes were sequenced. B) Timeline of the implementation of ITN (yellow, control) and ITN-PPF bednets (blue, treated) in each cluster, by year and month, for clusters where mosquitoes were sequenced. Numbers within a cell indicate the number of *An. gambiae* specimens that were sequenced in that particular cluster:timepoint with the total numbers per cluster given in the right hand column.

### Sequencing

A total of 16,785 mosquitoes were collected during this trial using CDC light traps, of which 14,489 (86%) were *An gambiae sensu strico* (hereafter *An. gambiae*). From these, we sequenced the whole genomes of 893 *An. gambiae*. We chose mosquitoes from clusters pre (ITN) and post-treatment (ITN-PPF) from around early (July/June 2014 and 2015), peak (September 2014 and 2015) and late (November 2014 and 2015) rainy season; while also ensuring we had good representation of space to capture the largest possible distances among clusters (**Figure 2**). To ensure sufficient sample sizes for population genetic analysis, we primarily restricted our choice to clusters x monthly timepoints for which >10 samples were available. The clusters selected were an average of 29km apart from each other (minimum of 3.53km, and maximum of 74.45km). In total we sequenced 240 *An. gambiae* specimens from ITN-PPF clusters, and 174 from ITN clusters in 2014, and 418 and 61 *An. gambiae* sampled from ITN-PPF and ITN clusters in 2015, respectively. Genomic DNA was extracted using a modified low salt Proteinase K buffer to maximise DNA yield^40^, and Illumina-sequenced by Novogene UK. Mean per-sample sequencing coverage was 3.43X (for per-sample sequencing data see **Table S1**). Samples were grouped for analysis by cluster, and by year/month of collection. Each sample grouping is hereafter referred to as a cluster:timepoint.

**Figure 2.**
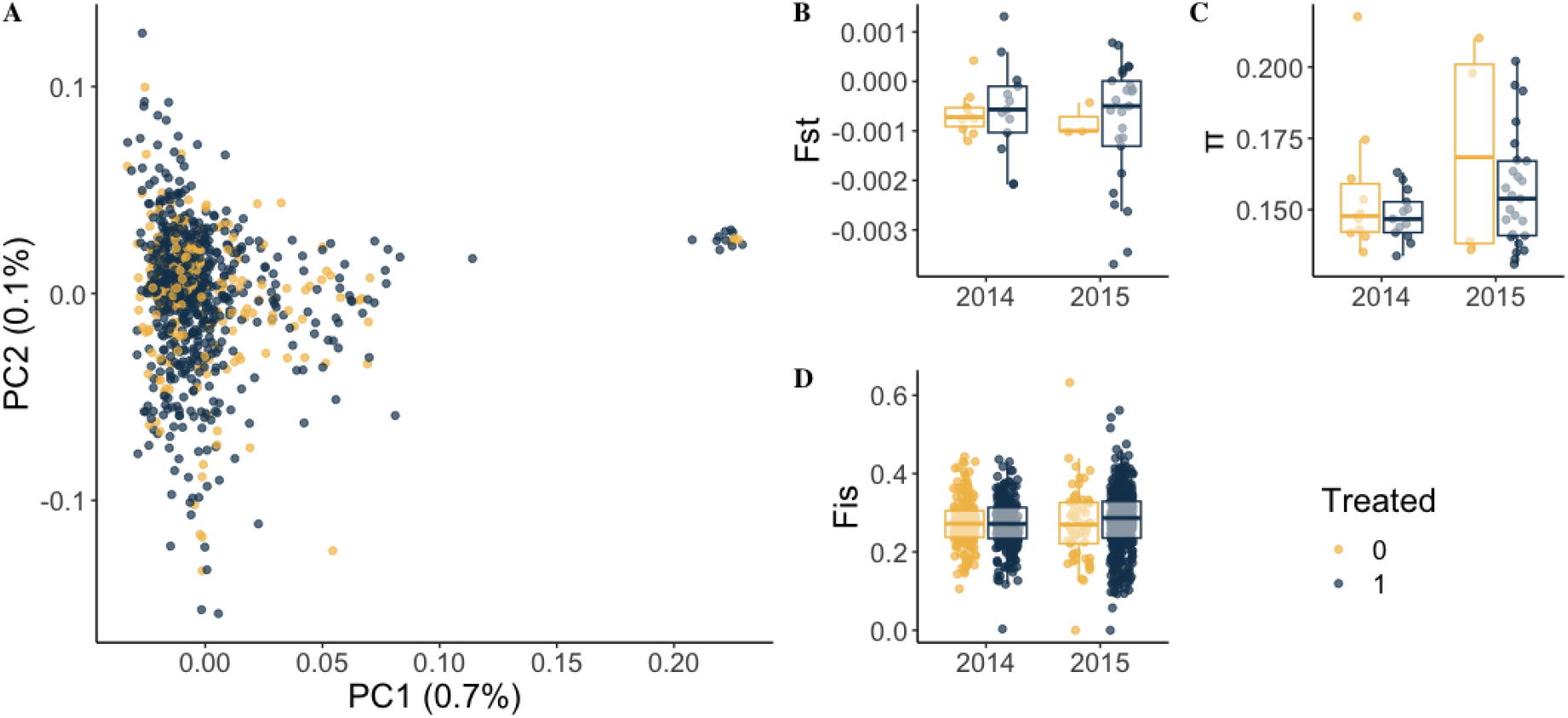
Analysis of spatial population structure and diversity. (A) Principal component analysis of *An. gambiae* samples, with variance explained by each PC in brackets. Panels B, C and D indicate between-cluster:timepoint *F*_*ST*_, within cluster:timepoint π, and per-individual *Fis* by treatment and year, respectively. Yellow and blue points indicate control (0) and treated (1) sites and individuals respectively.

### Population genetic analyses

We investigated the impact of ITN-PPF on the spatial genetic structure of *An. gambiae* during the study in multiple ways. First we estimated kinship between all mosquito samples with the aim of building a dispersal kernel from geographic distance between close kin^41^. This analysis aimed to determine whether kinship decayed significantly with geographic distance (i.e. displayed isolation-by-distance) and whether isolation by distance was greater between samples from treated/treated and treated/control vs control/control clusters^30,31^ (e.g. if dispersal occurred less frequently from treated sites). Second we estimated *F*_*ST*_ between groups of samples from each cluster:timepoint (year-month), and all other cluster:timepoints, to determine whether treated sites had higher mean genetic differentiation, and therefore potential population fragmentation^42,43^ (possibly due to reduced dispersal or diversity in smaller populations). Finally we performed a principal component analysis of genotype-likelihoods from all mosquito samples to test whether those collected from treated (ITN-PPF) clusters have more discrete genetic clusters than those from control (ITN) clusters^42,43^. We investigated the impact of the ITN-PPF intervention on the genetic diversity of *An. gambiae* using two metrics: nucleotide diversity (π), and individual inbreeding coefficient (*F*_*IS*_). We estimated *F*_*IS*_ per individual: an increase in per-individual inbreeding is a result of mating between related individuals, and of low genetic diversity, in each cluster:timepoint. Finally, we performed genome-wide scans of *F*_*ST*_ between treated and control sites to identify potential regions of the genome subject to selection in candidate insecticide-resistance regions^44^. The total sample size (893), and mean sequence coverage level (3.43), is sufficient to accurately estimate individual inbreeding coefficients^45^, kinship coefficients^46^, the site-frequency spectrum for the entire sequenced population (for PCA), and for treatment/year subdivisions (for *F*_*ST*_ between years and treatments for genome scans)^47^. Sequenced sample sizes within cluster:timepoints ranged from 3-20, with sample sizes <10 excluded from per-cluster:timepoint based analysis such as *F*_*ST*_, based on the expectation that low coverage sequencing of sample sizes between 10 and 20 are able to recover similar results as higher coverage and sample sizes^35^.

## Results

No discernible clustering pattern was present in an all-sample PCA (**Figure 2A**), which revealed two groups along PC1 (**Figure 2A**) but these did not correspond to any specific cluster, timepoint, or treatment arm. Instead, the small group beyond the value of 0.2 of PC1 are from mosquitoes of a mix of clusters and timepoints (**Figure S1, Figure S2**) indicating possible cryptic population structure associated with currently unknown factors. We were also not able to detect any spatial structure and consequently no change in spatial structure in mosquito populations in association with the intervention. The mean *F*_*ST*_ between a single cluster:timepoint, and all other cluster:timepoints, was very low (mean -0.00073), with a negative *F*_*ST*_ value denoting greater genetic diversity within populations than between them. We estimated slightly lower mean differentiation (*F*_*ST*_) between ITN-PPF clusters (−0.000748) compared to ITN clusters (−0.000680) and between 2015 (−0.000841) and 2014 (−0.000597) (**Figure 2B**), though neither difference was significant. These results suggest that the mosquito population did not detectably fragment or change in structure as a result of treatment as the trial progressed. We found no obvious potential dispersal events. Kinship coefficient values between 0.125 and 0.5 indicate close-kin relationships^44,48^. We found the mean kinship coefficient for our data was -0.66 [min= -2.95; max: -0.26] suggesting no close-kin relationships in any of the samples. Moreover, we found no significant change in between-sample relatedness in pre-or post-treatment clusters from 2014 or 2015.

The nucleotide diversity (π) of *An. gambiae* from ITN and ITN-PPF clusters in 2014 and 2015 was between 0.13 and 0.21, with a mean of 0.15, comparable to previous estimates of West African *An. gambiae*^33,49^. Nucleotide diversity was slightly lower in *An, gambiae* from treated vs control clusters in 2015 (**Figure 2C**) and inbreeding coefficient (*Fis*) slightly lower in individuals from ITN-PPF vs ITN clusters in 2014 (**Figure 2D**). However, in neither case was this statistically significant. Finally, a genome scan of *F*_*ST*_ between treated and control clusters did not show any evidence of peaks in regions containing insecticide resistance genes between ITN-PPF and ITN clusters (**Fig S3**), indicating a lack of differentiation in allele frequencies at these loci. Taken together, the results indicate that the previously estimated 22% reduction in mosquito population size caused by the ITN-PPF nets was not associated with alterations in the spatial genetic structure and diversity of *An. gambiae* in the trial region, or to produce signs of selection (linked to insecticide resistance, for example), during the time span of the trial.

## Discussion

During the AvecNet cRCT, the ITN-PPF nets produced a 12% reduction in clinical malaria in children, and a 22% reduction in mosquito density^38^ compared to pyrethroid-only ITNs. These epidemiological and entomological impacts did not coincide with detectable changes in mosquito genetic population structure between treated and control clusters as seen by our analysis of whole genome sequence data of *An. gambiae* mosquitoes, densely sampled over a fine spatial scale. Our findings suggest that an epidemiological benefit from a successful vector control intervention is not necessarily associated with detectable changes in the vector population itself. To our knowledge, this is the first study reporting population genetic data, coupled with corresponding entomological and epidemiological endpoints in a cRCT setting.

Depending on the magnitude and modes of action of the intervention, epidemiological impacts may arise in the absence of substantial changes in vector population size and structure. For example, small reductions in mosquito life span might not necessarily impact recruitment or population size but could be sufficient to reduce the entomological inoculation rate (EIR) and consequently human infection risk^50-52^. Similarly, interventions that work by changing mosquito behaviour (e.g. impeding their ability to find or feed on a host) might substantially reduce human exposure to biting without reducing vector population size. For example, spatial repellents that impede vector ability to locate or feed on a human host might increase the time required for mosquitoes to find a blood meal and their probability of becoming infected (fewer host contacts), but may not diminish mosquito population size if other non-human hosts are available to feed on (e.g. livestock). Such mechanisms reduce the EIR and cut malaria risk even in the absence of substantial vector population size reduction or local extinction. In the case of the AvecNet trial, the additional sterilising effect of pyriproxyfen was sufficient to reduce mosquito population sizes by 22% and the likelihood of clinical malaria in children by 12% in the ITN-PPF cohort without altering mosquito vector population structure or diversity.

Previously published work examining the impact of vector control on population genetic structure and diversity in concert with entomological or epidemiological indicators (albeit none in a cRCT setting)^42,53–55^, reveals no consistent pattern of entomological or epidemiological effect required to achieve a population genetic impact. In Dielmo, Senegal, ITN-associated reductions of malaria prevalence from 86% to 0.3% over 22 years, and near-elimination of *An. gambiae* in favour of *An. arabiensis*, was not accompanied by any changes in diversity or inbreeding^56^. By contrast, ITN administration in Papua New Guinea was associated with changes in *An. hinesorum* and *An. farauti* spatial population structure, as well as reductions of 38% in caught *Anopheles* and declines in malaria transmission intensity of 11.1 to 0.9%^42,53^. It is possible that the reduction in vector population density reported in the ITN-PPF clusters of the AvecNet trial, and any underlying transient impact on population allele frequencies, might not have been strong enough to be detectable in the face of homogenising gene flow. Even if ITN-PPF nets caused temporary local extinction of malaria vectors in some clusters, recolonisation through migration might occur quickly enough to be indistinguishable from population persistence. These data suggest using population genetic data as an indicator of a successful vector control outcome may be of limited use when the impact of vector control on entomological indicators is modest. This might also be true in some cases where an intervention elicits large epidemiological and entomological impacts, but large mosquito population sizes and immigrations from outside the trial area make it difficult to detect changes in in vector population genetic analyses. In the latter case, application of our approach to situations where evaluations are conducted at a larger spatial scale than a typical cRCT^57–59^ may have more utility.

We show that changes to population structure are not necessarily associated with significant epidemiological or entomological impacts of vector control, but population genetic techniques may become more reliably useful when the goal of an intervention is to eliminate malaria vector populations rather than just reducing human exposure. At low population sizes, acquiring adequate numbers of mosquitoes to robustly estimate population size may be challenging; whereas a small number of specimens may be sufficient to elucidate genomic signatures of vector population decline (e.g. inbreeding, differentiation). As discussions progress around the deployment of gene-drive technology, as well as other biological control tools that may eliminate mosquito vector populations ^60,61^, it is also likely that genomics-based approaches such as those developed here for estimating mosquito population diversity, structure and sizes will become crucial for measuring the impact of vector control.

## Acknowledgements

We would like to thank all teams, local communities and community leaders involved in the original AvecNet Trial. This research was funded by the European Research Council, European Union’s Horizon 2020 Research and Innovation Programme grant 852957.

## Data availability

Analysis code and bioinformatic workflows will be made available as a GitHub repository upon publication, and are uploaded as a supporting file to the journal. Raw reads are deposited in the European Nucleotide Archive under accession PRJEB62110.

## Author contributions

MV, TPWD, and DW conceived the ideas and designed methodology; WMH, N’FS coordinated and conducted the entomological analysis in the original trial and contributed the samples, SWL, N’FS, HR and AT conceived and designed the original trial and provided conceptual contributions, HMF and AS contributed irreplaceable conceptual support, PP performed DNA extractions, TPWD analysed the data, TPWD and MV led the writing of the manuscript. All authors contributed critically to the drafts and gave final approval for publication.

## Supplementary Material

### Supplementary Methods

We sequenced the whole genomes of 893 *An. gambiae s*.*s*. to a median depth-of-coverage of 3.58X with 2×150bp paired-end Illumina reads. Raw reads were trimmed using fastp^62^, and aligned to the *An. gambiae* PEST reference genome^63^, using bwa-mem^64^. Alignments were sorted, with duplicates marked, using samtools^65^. BAM alignments were grouped into cohorts for analysis, consisting of all 893 samples, and samples grouped by cluster and timepoint (year-month, see **Figure 1** for cohort sizes). Genotype-likelihoods (GLs) and site allele frequencies (SAFs) were inferred using ANGSD^66^. All-sample GL calling was performed for principal-component analysis (PCA) and inbreeding coefficient (Fis) estimation. Per-cluster per-timepoint site allele frequency estimation (see **Figure 1 for cohort sizes**) was performed on the whole genome for the estimation of per-cluster per timepoint *F*_*ST*_ and π. For the all-sample callsets, GLs were called with the reference genome as the major allele (−doMajorMinor 4), filtering GLs based on a a minor allele (maf) frequency of 0.01 (−minmaf 0.01), maximum SNP p value 0.05 (−snp pval 0.05), covered only by properly paired reads (−only proper pairs 1), and uniquely mapped reads with a mapping quality of 20 (−uniqueOnly 1 -minMapQ 20). For the per-cluster per-timepoint SAF estimation, we used the parameters above, but omitted depth, SNP p-value, and maf filters. Filtering that biases the SFS towards more frequent sites can substantially distort estimates of *F*_*ST*_ and π. Full parameters are detailed in the avecnet_popgen repository detailed in the main text, under bin/angsd scripts. Principal component analysis and per-sample Fis estimation was performed on chromosome arm 3L GLs using PCAngsd^67^. Folded site frequency spectra (SFS) were calculated from SAF files for all samples, as well as per-cluster and per-timepoint, using realSFS^66^. For estimating differentiation of each cluster and timepoint with respect to the wider metapopulation, two-dimensional(2d)-SFS were generated between the allsample SFS, and per-cluster per-timepoint SFS. Each 2d-SFS was used to infer Hudson’s *F*_*ST*_ between each pair of cluster:timepoints. Hudson’s *F*_*ST*_ was chosen as it is more robust to sample size differences between populations^68^. Per-cluster/timepoint SFS were used to estimate per-population π. PCA, *F*_*ST*_, π, and *F*_*IS*_ data were analysed in R v4.2.2^69^. GLMs assessing the effects of treatment and year on *F*_*ST*_, π, and Fis were performed using glm. Negative binomial GLMMs were performed using lme4^70^. Residuals were examined using DHARMa v0.4.6^71^. Plotting was performed with ggplot2^72^ library. To assess the effect of DUO on per-individual *F*_*IS*_, and per-cluster-, per-timepoint *F*_*ST*_, and π, Gaussian GLMs using the three parameters prior as a function of DUO administration and time, were parameterised. All three GLMs returned insignificant results. *Supplementary Figures*

**Figure S1.**
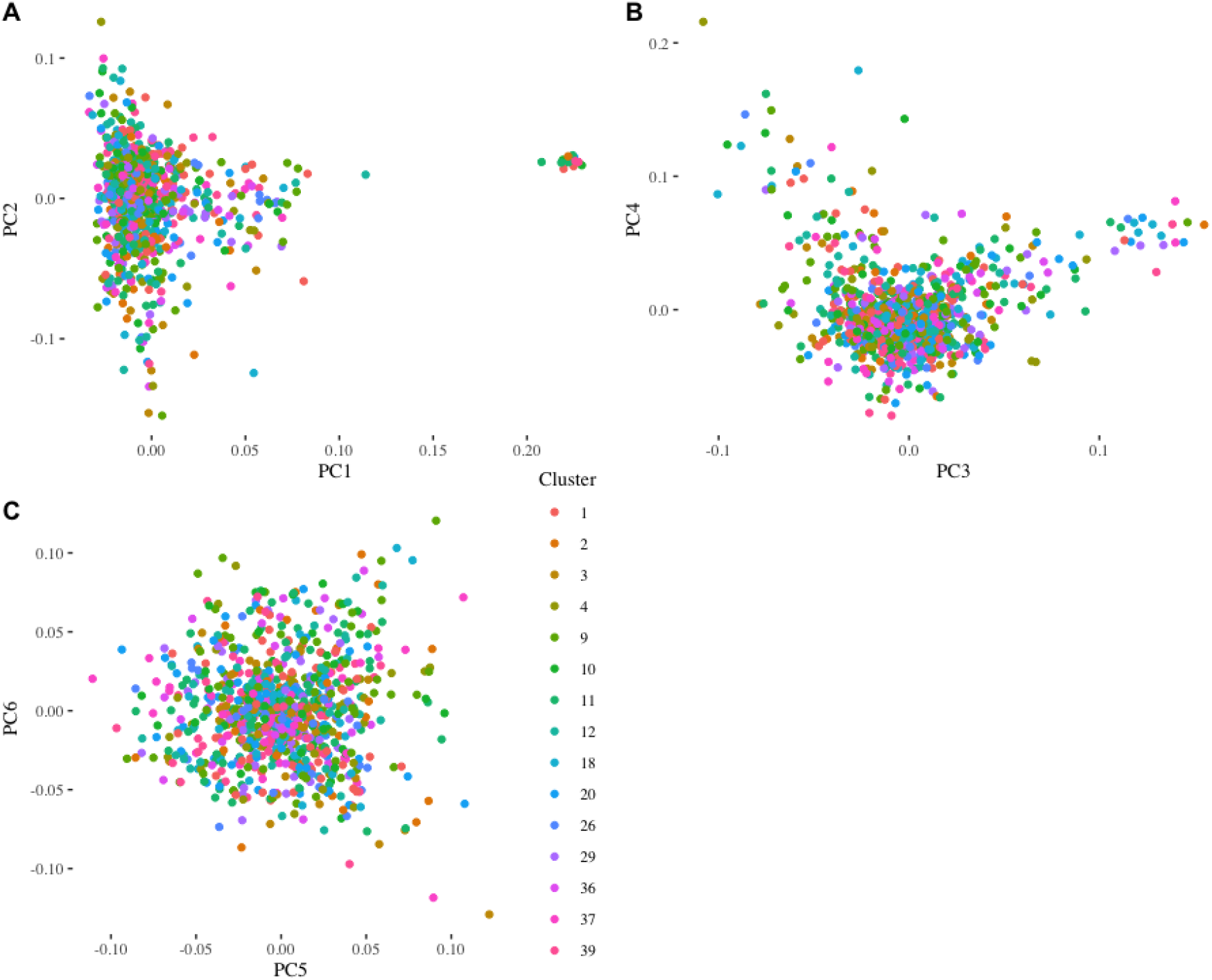
Principal component analysis (PCA) of sequenced *An. gambiae* from the AvecNet trial. Panels A, B and C indicate PCs 1/2, 3/4, and 5/6 respectively, indicated on the X and Y axes. Point colour indicates cluster.

**Figure S2.**
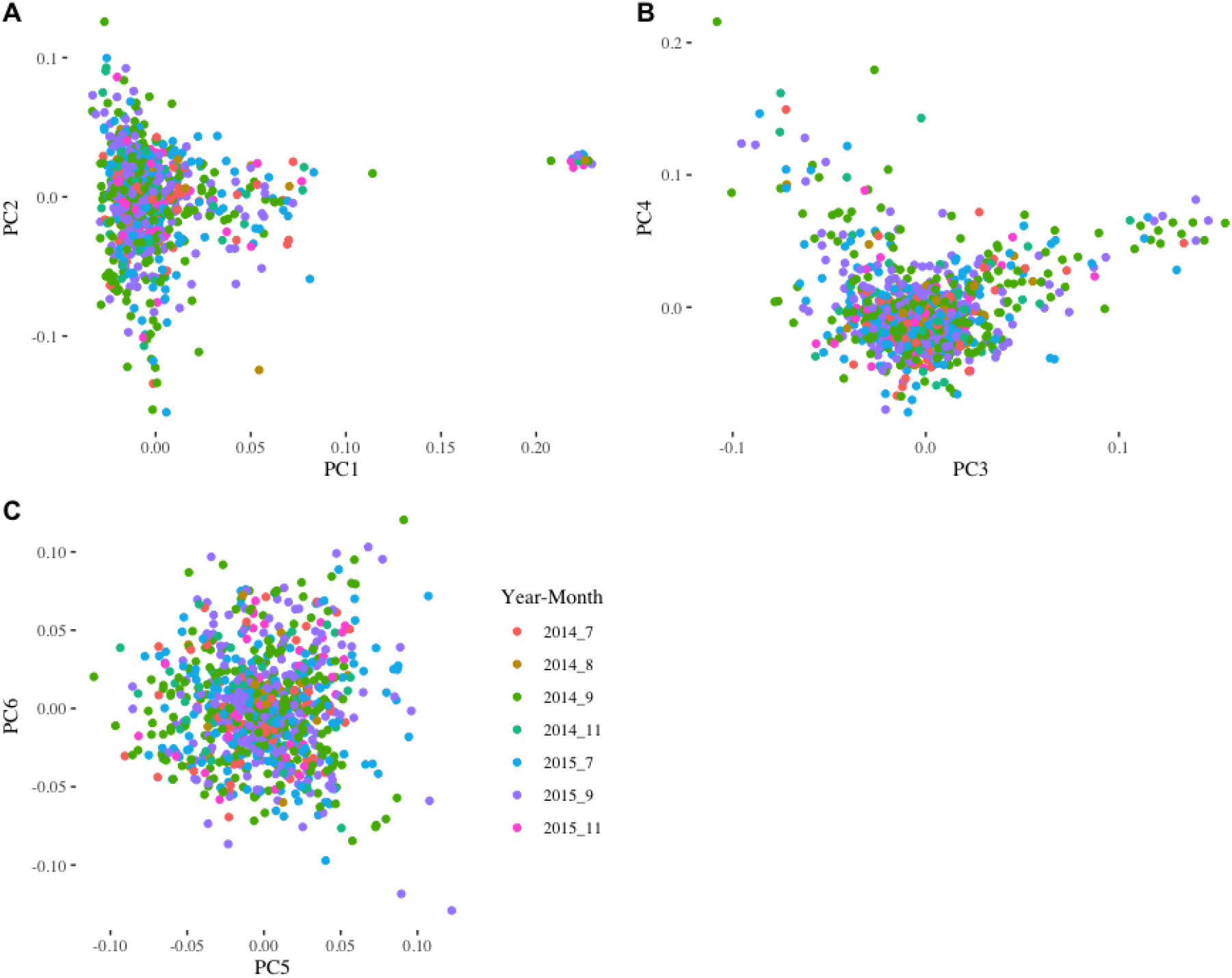
Principal component analysis (PCA) of sequenced *An. gambiae* from the AvecNet trial. Panels A, B and C indicate PCs 1/2, 3/4, and 5/6 respectively, indicated on the X and Y axes. Point colour indicates Year:Month.

**Figure S3.**
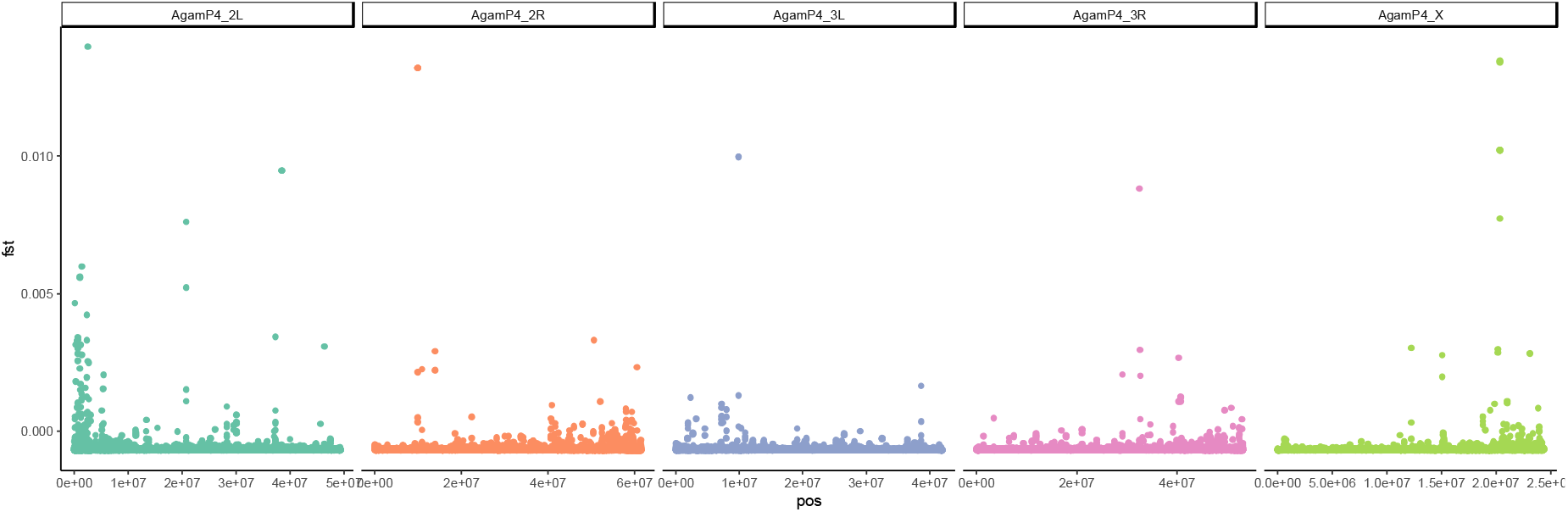
Genome-wide *Fst* between treated and untreated samples. X axis indicates chromosomal position, Y axis indicates *Fst* in 50Kb windows. Plot is faceted by chromosome

